# A novel mass assay to measure phosphatidylinositol-5-phosphate from cells and tissues

**DOI:** 10.1101/707604

**Authors:** Avishek Ghosh, Sanjeev Sharma, Dhananjay Shinde, Visvanathan Ramya, Padinjat Raghu

## Abstract

Phosphatidylinositol-5-phosphate (PI5P) is a low abundance lipid proposed to have functions in cell migration, DNA damage responses, receptor trafficking and insulin signalling in metazoans. However, studies of PI5P function are limited by the lack of scalable techniques to quantify its level from cells and tissues in multicellular organisms. Currently, PI5P measurement requires the use of radionuclide labelling approaches that are not easily applicable in tissues or *in vivo* samples. In this study, we describe a simple and reliable, non-radioactive mass assay to measure total PI5P levels from cells and tissues of *Drosophila*, a genetically tractable multicellular model. We use ^18^O-ATP to label PI5P from tissue extracts while converting it into PI(4,5)P_2_ using an *in vitro* kinase reaction. The product of this reaction can be selectively detected and quantified with high sensitivity using a liquid chromatography-tandem mass spectrometry platform. Further, using this method, we capture and quantify the unique acyl chain composition of PI5P from *Drosophila* cells and tissues. Finally, we demonstrate the use of this technique to quantify elevations in PI5P levels, from both *Drosophila* larval tissues and cultured cells depleted of phosphatidylinositol 5 phosphate 4-kinase (PIP4K), that metabolizes PI5P into PI(4,5)P_2_ thus regulating its levels. Thus, we demonstrate the potential of our method to quantify PI5P levels with high sensitivity levels from cells and tissues of multicellular organisms thus accelerating understanding of PI5P functions *in vivo.*

## Introduction

Phosphoinositides are a quantitatively minor class of glycerophospholipids, that mediate several cell biological functions that in turn can affect a wide range of physiological process. Phosphoinositides are generated by the selective phosphorylation of positions 3, 4 and 5 on the inositol headgroup of phosphatidylinositol [1]. The seven naturally occurring phosphoinositides are present in different amounts in the cell and each carries out distinct and characteristic functions within cells. Of these, the most recent phosphoinositide to be discovered was phosphatidylinositol 5-phosphate (PI5P) [2]. Since, its discovery, several studies have indicated that PI5P is present at low quantities in cells and its levels change in response to external cues such as UV radiation, oxidative or osmotic stress and growth factor stimulation [3-6] and such PI5P mediated signals might regulate cellular and physiological processes in multicellular organisms. Therefore, identifying and studying enzymes that can regulate PI5P levels in higher organisms is an active area of research.

Studies with cultured mammalian cells have shown robust changes in PI5P levels upon perturbations of two classes of phosphoinositide phosphate kinase (PIP kinase) enzymes; phosphatidylinositol 5 phosphate 4-kinase (PIP4K) and phosphatidylinositol 3 phosphate 5-kinase (PIKFYVE). PIP4K enzymes can phosphorylate PI5P to generate phosphatidylinositol 4,5 bisphosphate PI(4,5)P_2_and thus reduce PI5P levels in cells. On the other hand, PIKFYVE can synthesize PI5P directly by phosphorylating PI on the 5^th^ position of the inositol sugar ring [7]. Alternatively, PIKFYVE can phosphorylate PI3P to produce PI(3,5)P_2_ which can get dephosphorylated by a 3-phosphatase to form PI5P. However, the *in vivo* identity of such a 3-phosphatase is still elusive [8], Therefore, studying changes in PI5P levels from multicellular biological models where one or multiple PI5P regulating enzymes are manipulated, will develop a mechanistic understanding of PI5P under physiological conditions.

The quantification of phosphoinositides is typically done by one of two methods. The first involves the use of genetically encoded fluorescently tagged lipid binding domains [9]. This technique allows measurement of individual lipids that bind specifically to a protein domain at the level of a single cell with subcellular spatial resolution. In the context of PI5P quantification, the plant homeo domain (PHD) of the mammalian transcription factor, ING2 has been used in many studies [10,11]. However, due to its non-specific affinity toward PI3P, it is not regarded as an ideal probe for PI5P measurements [12].

A second approach is based on the detection and quantification of PI5P by radiolabelling cells with radioactive ^32^P ATP or ^3^H *myo*-inositol and then separating the deacylated monophosphoinositide isomers by ion exchange chromatography [13]. While this is a powerful approach in cultured cells, practical considerations restrict its use in animals, thus reducing the scope of its applicability for *in vivo* analysis. Some studies have used reverse phase HPLC to separate unlabelled deacylated PIP species and detect them by mass spectrometry [14,15]. However, reproducible separation of PI5P from the far more abundant and closely migrating PI4P is a challenge. More recently, various groups working on PI5P, have adopted a radioactive mass assay to measure PI5P levels [16,17]. The radioactive PljP-mass assay involves conversion of PI5P to PI(4,5)P_2_ by purified PIP4K α using an *in vitro* reaction that uses ATP with a ^32^P-label on its γ-P0_4_^3−^[^32^P ATP], This enables selective visualisation of the ^32^P labelled PI(4,5)P_2_ on a TLC plate [16]. While this technique is robust and offers good reproducibility, the disadvantage lies in the need to use radioactivity precluding the ability to handle a large number of samples at a given time and requires appropriate radiation safety facilities. A non-radioactive mass spec-based assay system, if available, can provide the advantage of avoiding potentially hazardous radiation and simultaneously offer higher sensitivity. To achieve these specific aims, we evolved the existing mass assay for PI5P levels to use a heavy Oxygen labelled ATP (^18^O-ATP) instead of using ^32^P-ATP in the kinase reaction. ^18^O is a non-radioactive stable heavy isotope of oxygen with 2 Da difference in mass from naturally occurring ^l6^O. This difference in mass allowed us to selectively monitor ^18^O-PI(4,5)P_2_ formed from biochemical PI5P by PIP4Ka, from a lipid mixture containing endogenous PI(4,5)P_2_ through the use of a liquid chromatography-tandem mass spectrometry (LC-MS/MS) based approach. In this study, we have developed a method based on this strategy to detect and measure changes in PI5P levels. Further, using the advantages of triple - quadrupole mass spectrometry, we were able to determine the levels of multiple species of PI5P, each with a unique fatty acyl chain composition.

## Materials and methods

### Fly strains and stocks

All experiments were performed with *Drosophila melanogaster* (hereafter referred to as *Drosophila).* Cultures were reared on standard medium containing corn flour, sugar, yeast powder and agar along with antibacterial and antifungal agents. Genetic crosses were set up with Gal4 background strains and maintained at 25°C and 50% relative humidity [18]. There was no internal illumination within the incubator and the larvae of the correct genotype was selected at the 3^rd^ instar wandering stage using morphological criteria. *Drosophila* strains used were ROR (wild type strain), *CIPIP4K*^*29*^ (homozygous null mutant of dPIP4.K), daGl4.

### S2R+ cells: culturing and dsRNA treatment

*Drosophila* S2R+ cells were cultured and maintained as mentioned in Gupta et. al., 2013 [19]. dsRNA treatment was performed as described in Kumari et al.,2017 [20]. Briefly, 0.5 × 10^6^ cells were incubated with 3.75 μg of dsRNA for 96 hours as described in Worby *et. al*, 2003 [21].

### Western blotting

Five wandering 3^rd^ instar larvae were used for lysate preparation. They were washed in PBS and homogenised using clean plastic pestles in lysis buffer [50 mM Tris/Cl-pH 7.5, imM EDTA, 1mM EGTA, 1% Triton X-100, 5omM NaF, 0.27 M Sucrose, 0.1% (β-Mercaptoethanol and freshly added protease and phosphatase inhibitors (Roche)]. Lysates were kept on ice for 15 mins following which the carcass was pelleted at loooXg for 15 mins at 4°C. About 75% of the lysate was transferred to a fresh tube, 6X Laemelli buffer added and samples heated at 95°C for 5 mins. Lysates of S2R+ cells were prepared by first washing the cells twice in 1× PBS and lysed in the lysis buffer mentioned previously, Laemelli buffer was added and the sample heated at 95°C for 5 mins. Proteins were separated by SDS/PAGE and transferred onto nitrocellulose membrane using wet transfer. Membranes were blocked with 10% Blotto (in 0.1% Tween 20,1×TBS{TBST}) and primary antibody incubations were performed in 5% BSA in 0.1% TBST overnight at 4°C. Washes were done in TEST. Secondary antibody incubations were performed for 2 hour at room temperature after which, the membranes were visualised using ECL reagent (Biorad). Dilutions of antibodies used: 1:4000 for anti-tubulin (E7-C), (mouse) from DSHB and 1:1000 anti Actin (Rabbit) A5060 from Sigma and 1:1000 for anti-dPIP4K antibody (Rabbit) used was generated in the lab and described previously [19].

### Lipid standards

diCi6-Pl3P - Echelon P-3016; diCi6-Pl5P - Echelon P-5016; Avanti 850173 | rac-i6:o PI(5)P-d5 (Custom synthesised); 17: o 20: 4 PI3P - Avanti LM-1900; 17: o 20: 4 PI(4,5)P_2_ - Avanti LM-1904,

### Lipid isolation

All the lipid isolation and processing steps were adapted from Jones et. al., 2013 [16] and Clark et. al., 2011 [22]. <u>Larvae:</u> For each sample, five wandering 3^rd^ instar larvae were washed, dried on a tissue paper and transferred to 0.5 ml tubes (Precellys Bertin corp. KT03961-1-203.05) containing 200μl Phosphoinositide elution buffer [PEB: chloroform/methanol/2.4 M hydrochloric acid in a ratio of 250/500/200 (vol/vol/vol)]. A Bertinhomogenizer instrument, Precellys®24 (P000669-PR240-A) was used at 6ooorpm for 4 cycles with 30 secs rest time on ice. The homogenate was transferred to 75011] of PEB in a 1.5 ml Eppendorf and sonicated for 2 mins. We added either 10 or 35ng of 17:0 20:4 PI(4,5)P_2_ as internal standard (in methanol) for LC-MS/MS experiments. Further, 250μ;l chloroform and 250μl MS-grade water was added and vortexed for 2 mins. The contents were then centrifuged for 5 mins at loooXg to obtain clean phase separation. The lower organic phase was washed with equal volume of lower phase wash solution [LPWS: methanol/i M hydrochloric acid/chloroform in a ratio of 235/245/15 (vol/vol/vol)] and vortexed and phase separated. The organic phase thus obtained was dried in vacuum and stored at −20°C and processed further within 24 hours, <u>S2R+ cells</u>: Cells were dislodged from the dishes, transferred to 1.5 ml Eppendorf tubes and washed twice with 1X Tris-buffered saline (TBS) following which 950 μl of PEB was added to the tubes. The mixture was vortexed for 2 mins following which it was sonicated in a bath sonicator for 2 mins. The rest of the procedure was identical to that described for extraction of lipids from larvae (see above).

### Neomycin chromatography

Glyceryl glass (controlled pore) beads were purchased from Sigma (cat. no. GG3000-200) and charged with neomycin sulfate as previously published [16] or neomycin beads were purchased from Echelon Biosciences (cat. no. P-B999). Chromatography was performed with 1-2 mg bead equivalent in slurry form on a Rotospin instrument (Tarsons, India) using buffers as described in Jones et. al., 2013.

### Total Organic Phosphate measurement

500 (μl flow-through obtained from the phosphoinositide binding step of neomycin chromatography was used for the assay. The sample was heated till drying in a dry heat bath at 90°C in phosphate-free glass tubes (Cat# 14-962-26F). The rest of the process was followed according to Jones et. al., 2013 [16].

### GST-PIP4K α based ^18^O-ATP mass assay

Either diCi6-Pl5P (Echelon) (for radioactivity) or 17:0 20:4 PI3P and d5-diCi6-Pl5P (for LC-MS) were mixed with 20 μM Phosphatidylserine (PS) (Sigma P5660) and dried in a centrifugal vacuum concentrator. For biological samples, the PS was added to the organic phase obtained at the end of the neomycin chromatography before drying. To this, 50 μl lomM Tris-HClpH 7.4 and 50 μl diethyl ether was added and the mixture was sonicated for 2 mins in a bath sonicator to form lipid micelles.

The tubes were centrifuged at loooXg to obtain a diethyl ether phase and vacuum centrifuged for 2 mins to evaporate out the diethyl ether. At this time, the reaction was incubated on ice for ca. 10 mins and 2X kinase assay buffer (100 mM Tris pH 7.4, 20 mM MgCl_2_, 140 mM KCl, and 2 mM EGTA and 1 μg equivalent GST-PIP4K enzyme-expressed and purified according to Jones et. al., 2013) was added. For the experiments with radioactivity, the 2X kinase assay buffer contained 40 μM cold ATP, 5 μCi [γ-^32^P] ATP (for synthetic lipids) or 10 μCi [γ-^32^P] ATP for biological lipids. For LC-MS/MS based experiments the kinase assay buffer contained 80 [γM ^18^O-ATP (OLM-7858-20, Cambridge Isotope Laboratory).

All the assays were performed at 30°C; in the case of synthetic lipids, assays were performed for defined periods of time; in the case of biological samples, assays were performed for 16 hours unless otherwise mentioned. Reactions were stopped by adding 125 γl 2.4N HCl, 250 γl methanol and 250 γl chloroform. The mixture was vortexed vigorously and spun down for 5 mins at loooXg to obtain clean phase separation. The lower organic phase was washed with equal volume of LPWS and vortexed and phase separated. The final organic phase obtained was processed further for either TLC or chemical derivatization to analyse the products of the reactions.

### Derivatization and LC-MS/MS

The organic phase obtained after lipid extraction was directly subjected to derivatization using 2 M TMS-diazomethane (Acros AC385330050), with all necessary cautions as mentioned in Sharma et al., 2019 [23]. After this, 50 γl TMS-diazomethane was added to each tube and vortexed gently for 10 min. The reaction was neutralized using 10 γl of glacial acetic acid. This was followed by two post derivatization washes as described in Sharma et al., 2019 [23]. To this final extract, 90% (v/v) methanol was added to this and dried for ∼2 hours in a centrifugal vacuum concentrator at 300 rpm. Next, 170 γl of methanol was added to the dried sample after which it was ready for injection. Samples were run on a hybrid triple quadrupole mass spectrometer (Sciex 6500 Q-Trap) connected to a Waters Acquity UPLC I class system. Separation was performed either on a ACQUITY UPLC Protein BEH C4, 300Å, 1.7 μm, 1 mm X 100 mm column [Product #186005590] or a 1 mm X 50 mm column [Product #186005589], using a 45% to 100% acetonitrile in water (with 0.1% formic acid) gradient over 10 mins or 4 mins. Detailed MS/MS and LC conditions are presented in tabular format in the next section.

### LC conditions and Mass Spectrometry parameters

#### Gradient conditions

| Time (in mins)<br>1 mm X 100 mm column | Time (in mins)<br>1 mm X 50 mm column | Flow rate<br>(ml/min) | % solvent A | % solvent B |
| --- | --- | --- | --- | --- |
| Initial | Initial | 0.100 | 55 | 45 |
| 5 | 0.50 | 0.100 | 55 | 45 |
| 10 | 4.50 | 0.100 | 0 | 100 |
| 15 | 7.20 | 0.100 | 0 | 100 |
| 16 | 7.30 | 0.100 | 55 | 45 |
| 20 | 10 | 0.100 | 55 | 45 |

##### MRM and NI. scans

Dwell time of 65 milliseconds were used for experiments with CAD value of - 2, GSi and GS2 at 20, CUR (Curtain gas) at 37, IS (ESI Voltage) as 5200 and TEM (Source Temperature) as 5200. The following table lists the distinct parameters used for all PIP species and PIP_2_ species.

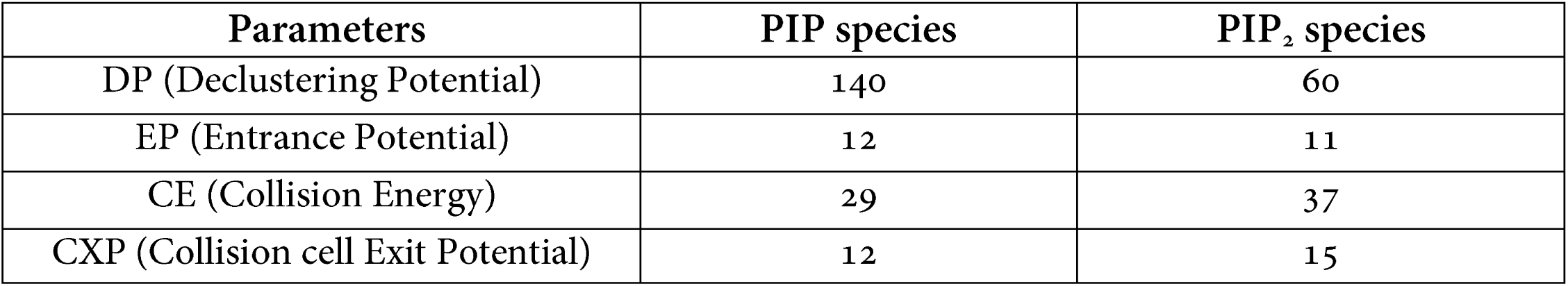

### Thin layer Chromatography

Extracted lipids were resuspended in chloroform and resolved by TLC (preactivated by heating at 90°C for 1 hour) with a running solvent (45:35:8:2 chloroform: methanol: water:25% ammonia). Plates were air dried and imaged on a Typhoon Variable Mode Imager (Amersham Biosciences).

### Software and data analysis

Image analysis was performed by Fiji software (Open source). Mass spec data was acquired on Analyst® 1.6.2 software followed by data processing and visualisation using MultiQuant™ 3.0.1 software and PeakView® Version 2.0., respectively. Chemical structures were drawn with ChemDraw® Version 16.0.1.4. Illustrations were created with BioRender.com. All datasets were statistically analysed using MS-Excel (Office 2016).

## Results

### ^18^O-ATP can be used in an *in vitro* PIP4K assay

Lipid extracts from biological samples are expected to contain a mixture of all the phosphoinositides including PI5P and PI(4,5)P_2_. As a result, any assay that measures PI5P in a mixture by converting it to PI(4,5)P_2_ must be capable of producing unique PI(4,5)P_2_ species distinguishable from the endogenous PI(4,5)P_2_ already present. To this end, we established a simple but innovative modification to the existing radioactive PI5P mass assay where we substituted the use ^32^P ATP to γ-P0_4_^3- 18^O labelled ATP [^18^O ATP]; a schematic of the reaction is shown in Figure 1A. This allowed us to detect and quantify the product generated in the reaction by liquid chromatography coupled to a tandem mass spectrometer (LC-MS/MS). We expressed and purified GST-PIP4Ka and tested its activity on 600 picomoles of synthetic deuterated PI5P (d5-diCi6-Pl5P) custom synthesized by Avanti Polar Lipids. Lipids were extracted at the end of the assay, and the contents derivatized using TMS-diazomethane and injected through an in-line C4 UPLC column connected to a triple quadrupole mass spectrometer. Multiple reaction monitoring (MRM) method was used to selectively follow the elution of individual lipids. This method detects and allows quantification of a signature parent-daughter ion pair for a given molecule with high sensitivity [24]. Fragmentation of any PIP and PIP_2_ parent ions results in a loss of a neutral head group of fixed masses of 382 and 490 Da respectively [25]. When ^18^O-ATP is used in this kinase reaction, both the mass of the parent PIP, product and consequently, the neutral fragment generated due to fragmentation, increases by 6 Da to 496 Da (Figure 1B indicates the ^18^O in red). The other major fragment generated is the charged diacylglycerol group whose mass depends on the length and saturation of the fatty acyl chain at *sn-* 1 and *sn-2* position and can be calculated theoretically (Figure 1B). Thus, the d5-PI5P molecule can be detected as the MRM parent/daughter ion transition of 938.5/556.5 (difference in mass: 382 Da), d5-PIP_2_ as 1046.5/556.5 (difference in mass: 490 Da) and d5-^18^O-PIP_2_ as 1052.5/556.5 (difference in mass: 496 Da). Using this rationale, we observed two major peaks from the reaction in which GST-PIP4Ka was incubated with d5-Pl5P (Figure 1C). The first peak at R_t_=9.84 min corresponded to a methylated ds-PIsP and the second peak at R_t_= 9.92 min corresponded to a methylated ds-^18^O PIP_2_ (Figure 1C). The response ratio, expressed as area under the curve of product (d5-^18^O-PIP_2_) divided by the substrate (PI5P) was around 1.9 showing that there was significant production of ds-^18^O-PIP_2_ in the reaction (Figure 1D). Interestingly, a very small amount of d5-PIP_2_ was obtained (response ratio ∼ 0.02) that could arise from the ^16^O-ATP impurity in the commercial ^18^O ATP (Figure 1C’).

**Figure 1:**
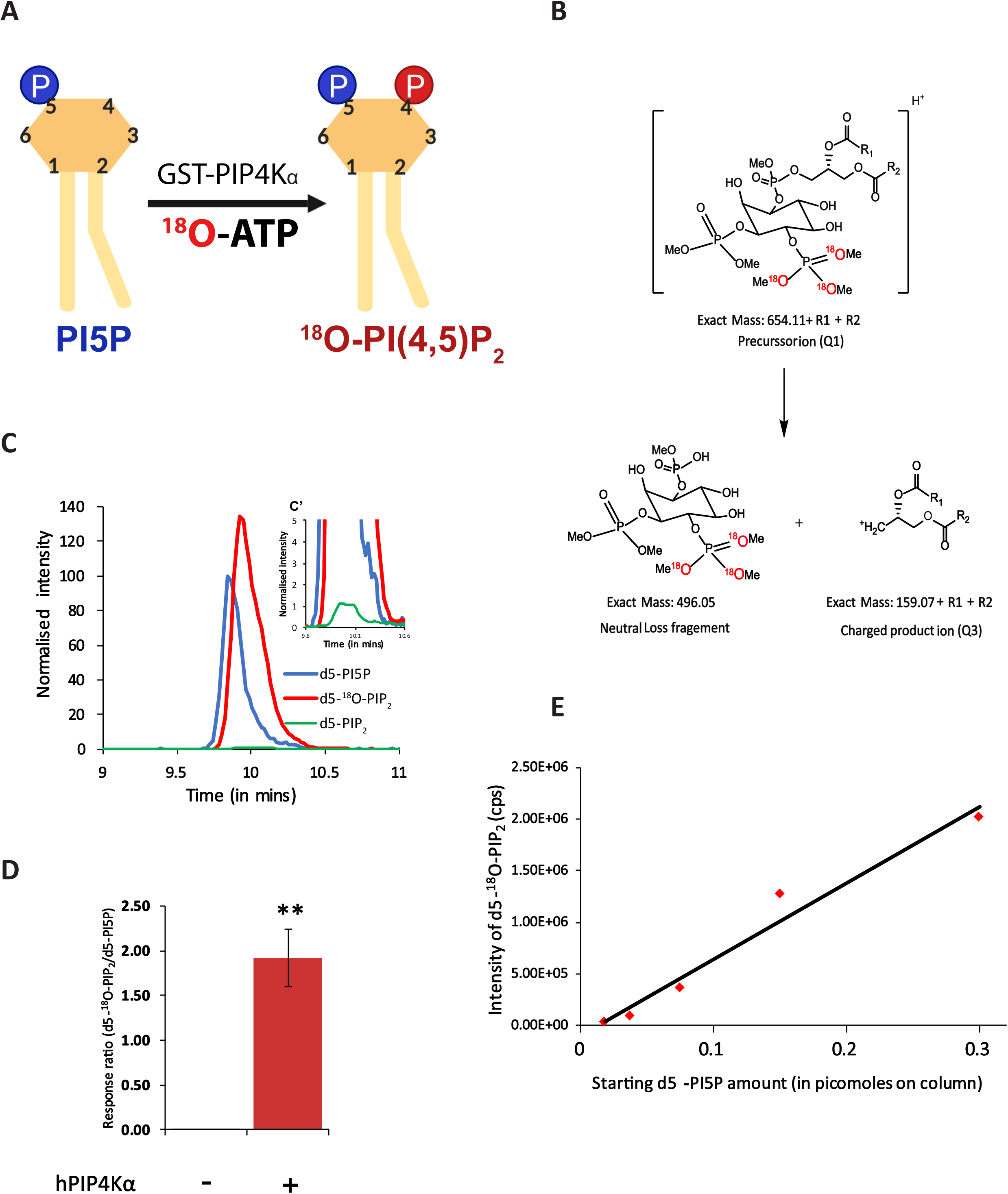
^18^O-ATP used as a substrate in an *in vitro* PIP4K assay. (A) A schematic to show the reaction catalysed by GST-PIP4Ka for a kinase reaction on PI5P (see methods for details) (B) Diagram representing methylated ^18^O-PI(4,5)P_2_ with +1 charge and the resultant products after fragmentation yielding a neutral head group of mass 496 Da and a charged diacylglycerol (DAG) fragment of mass = Parent mass (PM) - neutral loss (NL) mass. (C) Chromatogram representing the extracted ion chromatogram (XIC) of MRM transitions of substrate (ds-PIsP) in blue (938.5/556.5), the ^18^O-d5-PIP_2_product in red (1052.5/556.5) and d5-PIP_2_ product in green (1046.5/556.5). C’ shows that the ds-PIP, product formed in the reaction is negligible (∼ 1%). (D) Quantification of response ratio of d5-^18^O-PI(4,5)P_2_ formed from the earlier chromatogram. Number of individual reactions performed (n) = 3. Student’s unpaired t-test with 95% confidence interval shows significance between the no enzyme (bar 1) and with enzyme (bar 2). (D) A dose response curve of d5-Pl5P ranging from 0.01 picomoles to 0.3 picomoles on column. Y-axis depicts intensity of d5-^18^O-PI(4,5)P_2_ (in cps) and X-axis represents the amount of d5-PI5P loaded on column.

Usually, when working with biological samples, radioactive mass assays have reported a minimum detection limit of 1 picomole of converted PI5P [16]. We observe that in our LC-MS/MS method, we could linearly detect the conversion of PI5P to ^18^O-PIP_2_, even at a sub-picomole range when these lipids were loaded on column (Figure 1E). The conversion of such low amounts of PI5P was possible only with 16 hours of incubation and not with a l-hour incubation (data not shown). Based on this observation, we decided to conduct our assays with biological samples, where PI5P levels are low, for 16 hours.

### GST-PIP4K α is suitable for assaying PI5P by LC-MS/MS method

It is known from previous studies that in addition to PI5P, as a substrate, PIP4KS have a lower but significant *in vitro* activity on PI3P [26]. The radioactivity based PI5P mass assay can distinguish between the products formed from these two substrates because of the difference in migration of labelled PI(4,5)P_2_ and PI(3,4)P_2_ on a one dimensional TLC (Figure 2A). However, since there is presently no robust method to distinguish intact PIP_2_ isomers on LC-MS platforms, it was prudent to expect that the total PIP_2_ formed in the PIP4Kα kinase assay performed on lipids extracted from biological samples could have some contribution from PI(3,4)P_2_ in addition to PI(4,5)P_2_. We performed control experiments to estimate the amount of PI(3,4)P_2_ that might be formed in our ^18^O-ATP based kinase assay. To estimate this, we calculated the slopes of product formation when using increasing concentrations of synthetic PI3P or PI5P. At first, using the radioactivity (^32^P ATP) based kinase assay, we observed that the intensities of PIP_2_ spots produced from diCi6-Pl5P and diCi6-PI3P substrate were very different over the same range of substrate concentrations (Figure 2A). The slope ofPI(4,5)P_2_ formation from PI5P was 22 times steeper than that of PI(3,4)P_2_ formation from PI3P over a nanomole concentration range of each substrate (Figure 2B). Lipid samples prepared from cells for the kinase assay will have a mixture of PI5P and PI3P, wherein the levels of PI3P would be equal or higher than that of PI5P by about 2-3 times [13,27].

**Figure 2:**
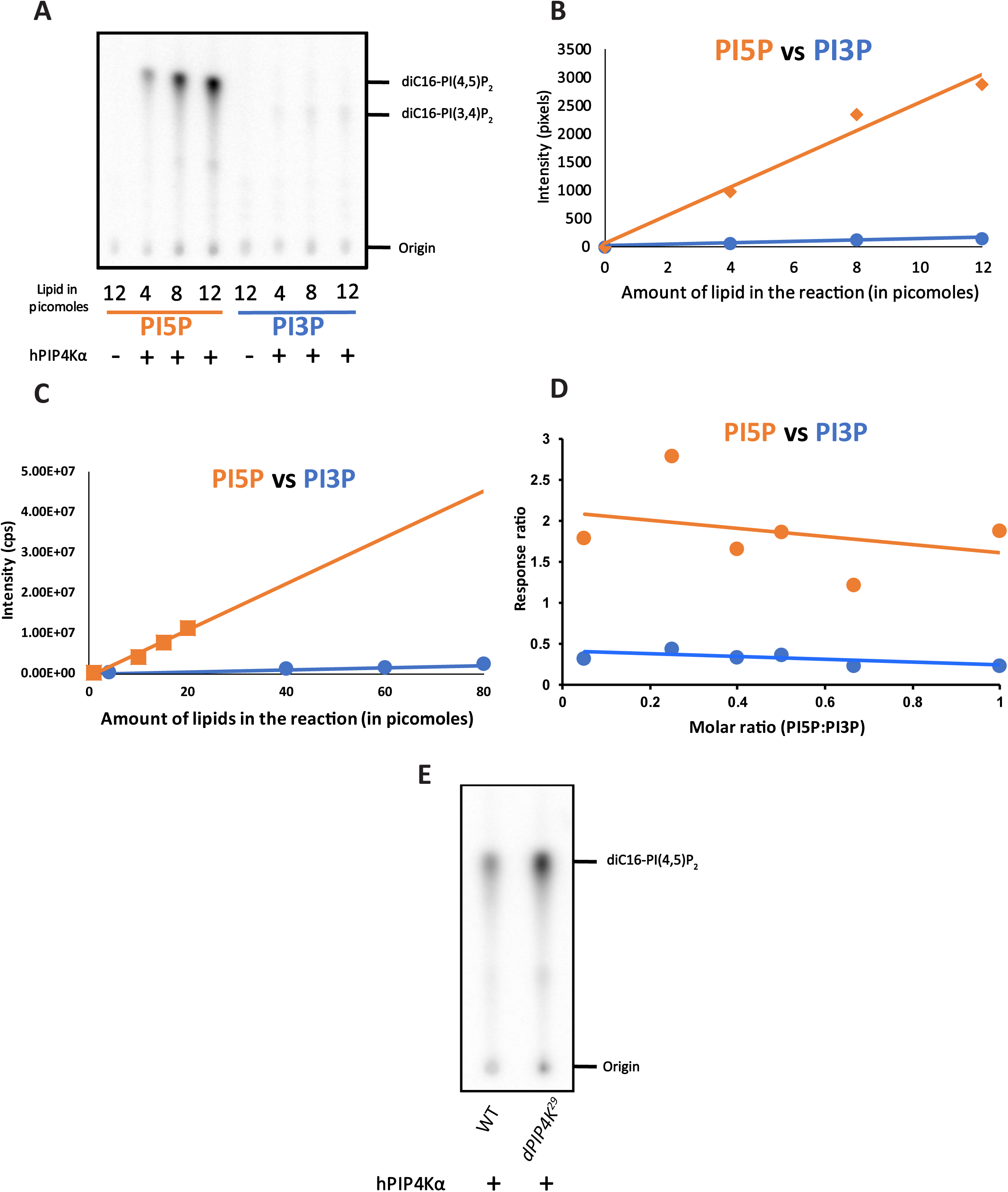
LC-MS/MS based assay reveals that GST-PIP4Kα is highly specific for PI5P and has lower affinity for PI3P. (A) TLC shows radioactive products formed from increasing concentrations of diCi6-Pl5P or diCi6-Pl3P (Echelon) upon kinase reaction with GST-PIP4Ka (see methods for details). Both assays have a ‘no enzyme control’ for 12 picomoles of lipid. PI(4,5)P_2_ or PI(3,4)P_2_ spots migrate with different R, and are labelled on the TLC along with the origin (B) Pixel intensity of the TLC image from (A) has been quantified and plotted for both PI5P and PI3P. Equations for PI5P: y = 250.32X + + 53-897; PI3P: y = 12.771X + 9.5691 (C) Intensity response (cps) of respective products of PI3P and PI5P, using different substrate amounts from a ^18^O ATP based kinase assay experiment analysed by LC-MS/MS using ds-diCi6-Pl5P or 17:0 20:4 PI3P as indicated in the X-axis. Equations for PI5P: y = 575245x - 841692; PI3P: y = 25265x - 39253 (D) Response ratio of PI5P vs PI3P at different molar ratios of PI5P to PI3P. Y-axis represents response ratio of either d5-^18^OPIP_2_to ds-PIsP (PI5P) or 17:0 20:4 ^18^O-PIP_2_to 17:0 20:4 PI3P and X-axis represents molar ratio of Pl5P:Pl3P (E) TLC shows single radioactive spot corresponding to PI(4,5)P_2_ from larvae of Wild type (WT) or *dPIP4K*^*29*^ (mutant of dPIP4K).

Next we tested the relative amounts of PI(3,4)P_2_ vs. PI(4,5)P_2_ formed by PIP4Kα when presented as a mixture of PI3P and PI5P. We used synthetic 17:0/20:4 odd chain PI3P (MRM: 995.5/613.5) and diCi6-d5-Pl5P (MRM: 938.5/556.5) which have distinctive masses despite being PIP isomers, thus allowing easy identification and quantification of the respective products formed from a mixture of PI3P and PI5P on a mass spectrometer. To test the extent of PI(3,4)P_2_ and PI(4,s)P_2_ formation from these two substrates, we mixed them at a molar ratio of 1:4, Pl5P:Pl3P, mimicking the relative abundances of these two lipids in cells. We observed that while there was detectable PI(3,4)P_2_ formation (MRM: 1109.5/613.5; difference of 496 Da) from the PI3P substrate (albeit at higher amounts of starting substrate), the dependence of PI(3,4)P_2_ formation on PI3P concentration had a lower slope compared to that for PI(4,5)P_2_ formation from PI5P (Figure 2C) and was similar to that observed in the radioactivity based PIP4Kα assays. Theoretically, in lipid extracts of genetic mutants of PI5P regulating enzymes, there will be a change in the molar ratios of PI5P to PI3P from the ratio in normal conditions. We observed that the response ratio of products formed from PI5P were always greater than compared to PI3P across a range of molar ratios of Pl5P:Pl3P, assayed under the given conditions, indicating that PIP4Kα will estimate PI5P much better over a broad range (Figure 2D). In summary, these data suggest that in our LC-MS/MS based assay (henceforth to be called the ^18^O-ATP mass assay), recombinant PIP4Kα can potentially be used to assay PI5P levels from a mixed phosphoinositide pool with limited contribution from PI3P.

A significant amount of PI(3,4)P_2_ was observed mainly when micromole amount of PI3P was used as substrate (Figure 2B). However, such high amounts of PI3P is unlikely to occur in small biological samples. To test this, we extracted total lipids from wandering 3^rd^ instar *Drosophila* larvae of wild type (WT) and compared it to *dPIP4K*^*29*^ mutant, where PI5P levels are higher. We noted a single spot of PI(4,5)P_2_for both samples on the TLC as reported earlier [16] (Figure 2E). We did not observe a separate spot on the TLC, corresponding to PI(3,4)P_2_ that migrated with a distinct mobility compared to PI(4,5)P_2_. This implied that the amount of PI3P used by GST-PIP4Kα, to generate PI(3,4)P_2_ under these assay conditions was negligible (Figure 2E).

### ^18^O-ATP based mass assay allows detection of multiple PI5P species in *Drosophila*

The strategy of methylating low abundance and poorly ionisable lipids such as PIP3 was initially described by Clark et.al [22]. If this approach is applied to the biological lipids extracted at the end of our ^18^O ATP mass assay, one should be able to detect low abundant methylated ^18^O-PIP_2_ products (precursor ion in Figure 1B). As described earlier, fragmentation of these precursor ions should generate (i) a neutral head group of fixed mass 496 Da and (ii) a charged diacylglycerol fragments whose mass would depend on the length and saturation of the fatty acyl chain at *sn-i* and *sn-2* positions (R_1_, and R_2_ in Figure 1B). After performing the ^18^O-ATP mass assay with lipid extracts from wild type larvae, we used neutral loss scanning (NLS) to identify all the precursor ions which generate fragments after loss of neutral mass 496 Da; this approach allowed us to identify eight such parent masses (Figure 3Ai) that produced a neutral loss fragment of 496 Da. This result implies the existence of eight molecular species of PI5P that were captured during the ^18^O-PIP_2_ assay. The acyl chain lengths of these ^18^O-PIP_2_ were calculated by subtracting 496 Da from the observed parent masses (Table1). Next, we performed a neutral loss scan for 382 Da corresponding to the neutral head group that could be generated from PIP species using the same lipid extract as above. The masses of ^18^O-PIP_2_ species detected correlated well with the masses of the corresponding PIP molecules (Figure 3 Aii). Using this information and theoretical calculation, we set up multiple reaction monitoring (MRM) methods with the Q1/Q3 masses mentioned in Table1 and detected the intensities of the various ^18^O-PIP_2_ species from wild type *Drosophila* larval samples (Figure 3B). Using this MRM method, we detected all eight individual species described above, thus confirming our results from the Neutral loss scans. We observed that the 34:2 species was the most abundant followed by 36:3, 36:2, 34:3 and 32:1. These data correlate well with the molecular species of PIP_3_ recently reported from *Drosophila* larval tissues [23]

**Figures 3:**
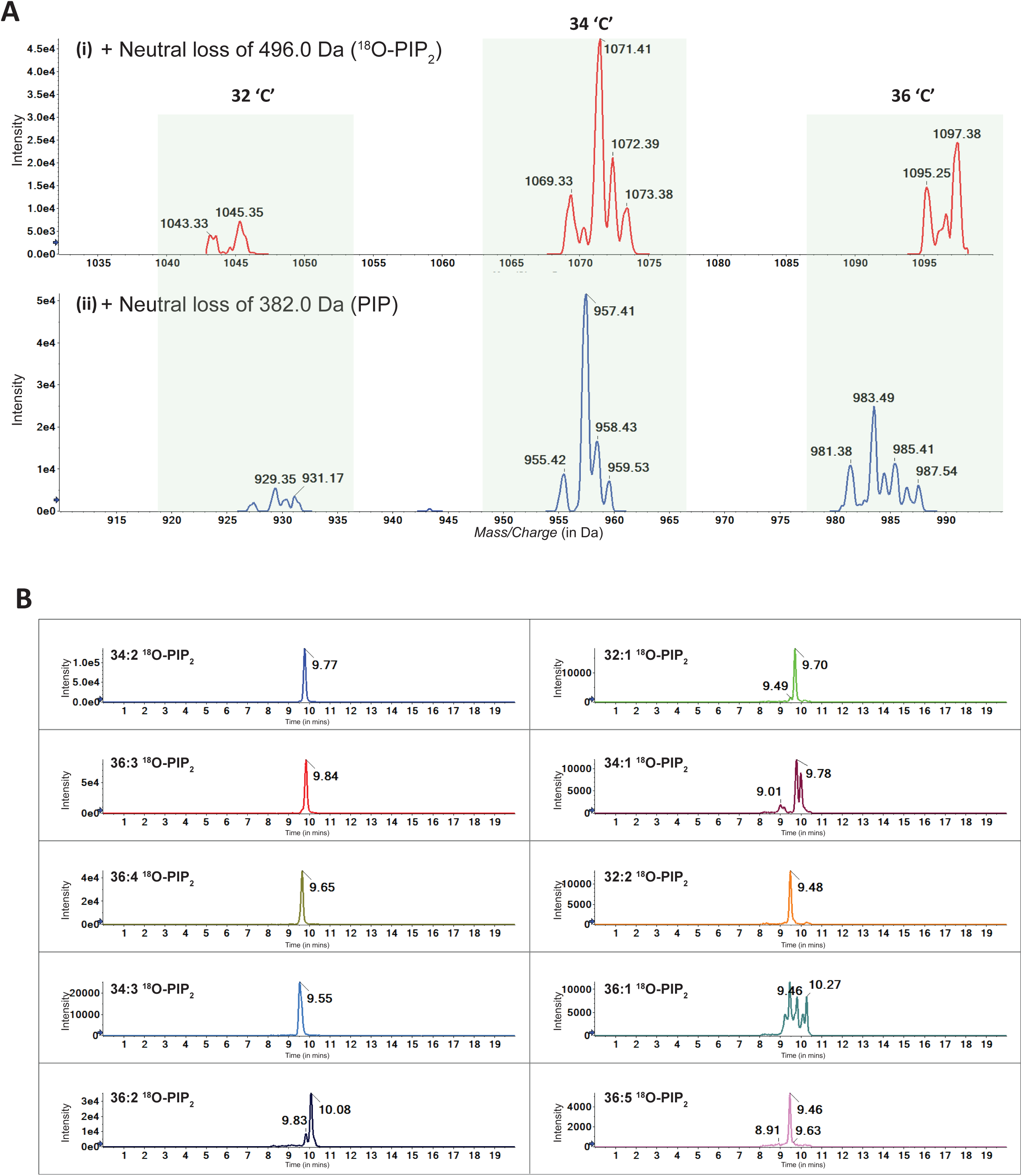
Mass spectrometry setup for ^18^O-ATP based mass assay of biological PI5P from *Drosophila*. (A) (i) Spectrum from + Neutral Loss of 496.0 Da (scanned for 1000 - 1245 Da) in a 20 mill run. (ii) Spectrum from + Neutral Loss of 382.0 Da (scanned for 750 - 1245 Da) in a 20 min run. The Y-axis indicates intensity of ions and the X-axis represents Mass/Charge (in Da). The peaks are marked by masses which feature as parent masses of the respective neutral loss mass. Both scans were performed from WT assayed larval samples. The highlighted regions depict parent lipid species with a total of 32 carbon (32 ‘C’), 34 carbon (34 ‘C’)_and 36 carbon (36 ‘C’) atoms in the acyl chains; ^18^O-PIP_2_ species are also tabulated in Table 1 (B) Extracted ion Chromatogram (XIC) of WT larval assayed samples for the species that were picked up in NL scans. For each of the chromatograms, the Y-axis represents intensity of the MRM corresponding to the species, and the X-axis represents time on the LC. The chromatograms are arranged in decreasing order of ion abundance.

### Loss of *Drosophila* dPIP4K results in elevated PI5P, measured as ^18^O-PIP_2_ signal

We tested the ability of our method to detect and quantify the increase in PI5P levels earlier reported from adult flies of *dPIP4K*^*29*^, a protein null allele of *Drosophila* PIP4K [19]. An immunoblot was performed to confirm that the mutant larvae did not have any dPIP4K protein (Figure 4A). Figure 4B describes the workflow used to estimate PI5P levels from biological sample. The 3^rd^ instar wandering larval stage of *Drosophila* is a good model to study changes in growth and development in the organism. Since *dPIP4K*^*29*^ larvae show defects in growth and cell size, studying the regulation of PI 5 P in such a system will allow us to understand the relevance of this lipid to cellular growth and metabolism. To this end, we compared PI5P levels between wild type and *dPIP4K*^*29*^*K* larvae. PI5P was measured using ^18^O-PIP_2_ levels relative to an internal standard, and normalised for tissue size by a total organic phosphate measurement obtained from the third step of the sample preparation (Figure 4B and methods). We found increased PI5P in *dPIP4K*^*29*^ consistent with previous observations using radioactive mass assay that reported that PI5P levels are elevated in *dPIP4K*^*29*^ [19]. Figure 4C captures the acyl chain length diversity of larval PI5P; all of the eight detected species of ^18^O-PIP_2_ were elevated in *dPIP4K*^*29*^ implying that the corresponding species of PI 5 P were elevated in *dPIP4K*^*29*^. In *Drosophila* larval extracts, the major species of PI5P are those with acyl chain 34:2 and 36:3; both of these were elevated. In addition, six other species of unique acyl chain length that we could detect were also elevated.

**Table 1:** List of MRMs The mass of each parent ion detected in Q1 representing ^18^O-PIP_2_ is shown in column-Parent ion mass (Q1). The singly protonated fragment corresponding to a diacylglycerol fragment derived from each such parent ion is shown in column-Daughter ion mass (Q3). The acyl chain composition of this diacylglycerol fragment is shown in column-Computed Acyl chain length.

| Parent ion mass (Q1)<br><sup>18</sup>O-PIP<sub>2</sub> mass | Daughter ion mass (Q3)<br>Diacylglycerol mass | Computed acyl chain<br>length |
| --- | --- | --- |
| 1043.5 | 547.5 | 32:0 |
| 1045.5 | 549.5 | 32:1 |
| 1069.5 | 573.5 | 34:0 |
| 1071.5 | 575.5 | 34:2 |
| 1073.5 | 577.5 | 34:1 |
| 1093.5 | 597.5 | 36:5 |
| 1095.5 | 599.5 | 36:4 |
| 1097.5 | 601.5 | 36:3 |
| 1099.5 | 603.5 | 36:2 |
| 1101.5 | 605.5 | 36:1 |

**Figure 4:**
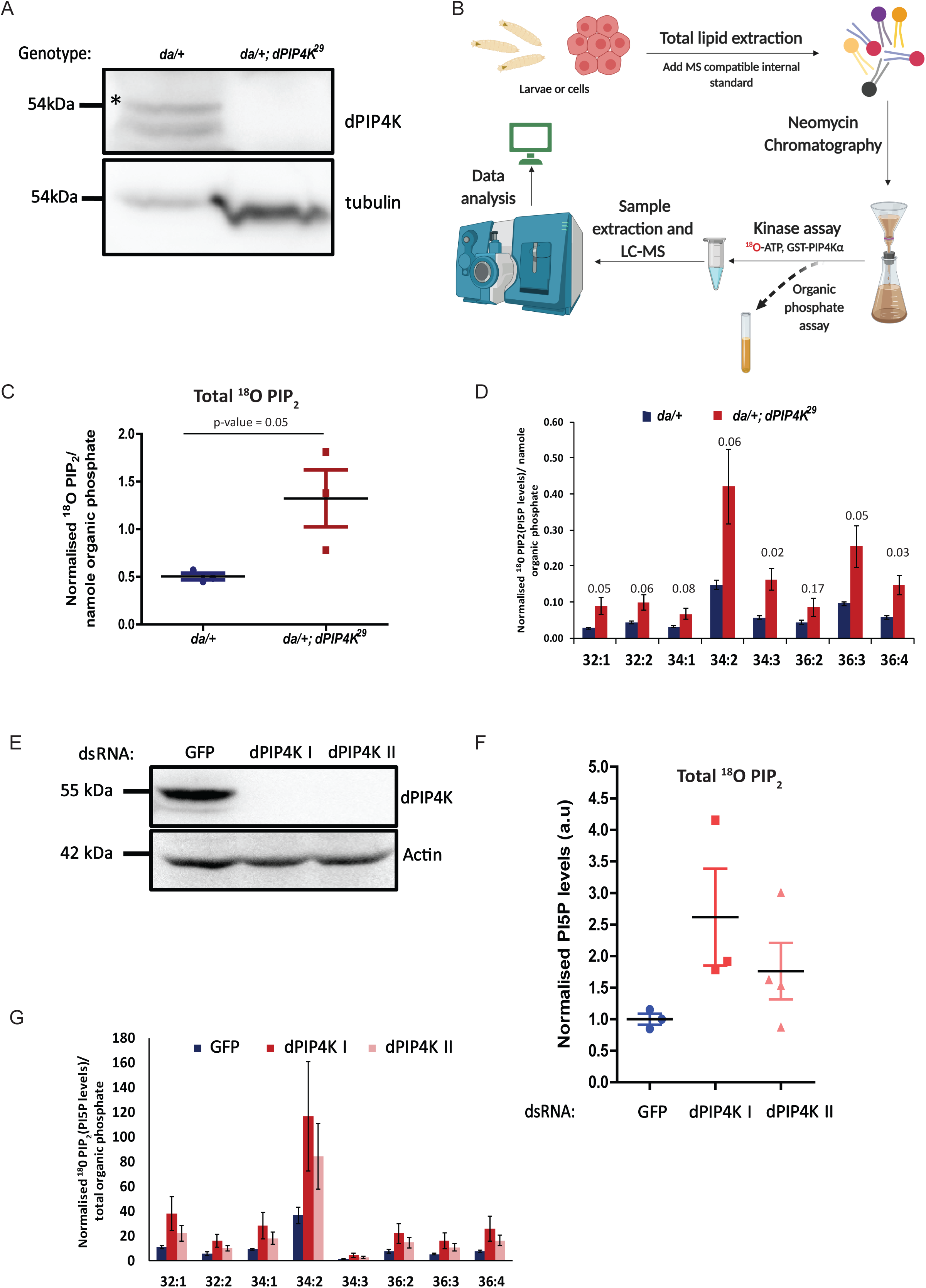
Loss of dPIP4K results in elevated in PI5P, measured as ^18^O-PIP_2_ levels in *Drosophila*. (A) Western blot showing dPIP4K protein expression in da/+ and da/+; *dPIP4K*^*29*^ samples prepared from 5 larvae. Tubulin is used as loading control. (B) Schematic summarising the methodology followed to perform the assay from *Drosophila*, larval or S2R+ cells (C) Total ^18^O-PIP_2_ normalised to internal standard (17:0 20:4 PI(4,5)P_2_) divided by organic phosphate value from processed *Drosophila* larval samples of da/+ (pan larval GaU control) and da/+; *dPIP4K*^*29*^ (dPIP4K mutant). n=3 where each sample has been prepared from five 3^rd^ instar wandering larvae (see methods for details). Error bars represent S.E.M. p value by student’s two tailed unpaired t-test is provided on the graph. (D) Different acyl chain length species of ^18^O-PIP_2_ of the data plotted in (B). Error bars represent S.E.M. Individual p value by student’s two tailed unpaired t-test is provided above each bar on the graph. (E) Western blot showing dPIP4K protein expression in S2R+ cells treated with dsRNAs against GFP (Control) or dPIP4K dsRNA I or dsRNA II. Actin is used as loading control. (F) Total ^18^O-PIP_2_ normalised to internal standard (17:0 20:4 PI(4,5)P_2_) divided by organic phosphate value from processed *Drosophila* samples of GFP, dPIP4K I or dPIP4K II dsRNA treated S2R+ cells. n=3 where each sample has been prepared from starting cell density of 0.5 million cells. The samples have been pooled across two days and have been normalised to the GFP dsRNA values (see methods for details). The graph has been normalised taking the GFP dsRNA value as 1. Error bars represent S.E.M. (G) Representative graph showing different acyl chain length species of ^18^O-PIP_2_ normalised to internal standard (17:0 20:4 PI(4,5)P_2_) divided by organic phosphate value from 2 samples of (F).

We also assayed PI5P levels from *Drosophila* S2R+ cells that had been depleted of dPIP4.K using with two different dsRNAs [recently reported in [23]]. Under our experimental conditions, treatment with either dsRNA resulted in complete depletion and dPIP4K protein could not be detected by western blot analysis (Figure 4E). The ^18^O-PIP_2_ mass assay using lipid extracts from these cells showed elevated levels of PI5P with a slight difference in the extent of PI5P elevation induced by each dsRNA (Figure 4F). We also estimated the level of each of the eight molecular species of PI5P and found that all of them were elevated on depletion of dPIP4K (Figure 4G). Thus, both in *Drosophila* larvae and S2R+ cells, using our ^18^O-ATP mass assay, we found that the major species of PI5P were elevated upon depletion of dPIP4K demonstrating that measurements done with our new method compare well with those previously reported using the radioactive mass assay.

## Discussion

20 years since the discoveiy of PI5P, the mechanisms by which its levels are regulated *in vivo* still remain unclear and is a matter of great interest owing to its proposed roles in various cell biological processes like cell growth, cell migration, autophagic and stress responses and apoptosis [28]. A key challenge in addressing this question has been the lack of a robust method to quantify PI5P. Till date, PI5P detection has relied principally on the use of analytical techniques that use radioactive labels [17] that are challenging to use on *in vivo* models. Here we describe a robust method to measure PI5P levels that does not require the use of radiolabelling and hence avoids associated hazards. The availability of our method offers additional advantages over the use of radiolabelling approaches. These include better control over the data variation due to the use of an internal standard for quantification in LC-MS/MS experiments and the ability to separately quantify each species of PIP and PIP_2_ separately. The use of this method in conjunction with genetically tractable metazoan models for *in vivo* analysis should facilitate the analysis of PI5P turnover *in vivo.* Our method, an adaptation of the conventional radiolabel based conversion of PI5P to PI(4,5)P_2_ shows superior sensitivity to the conventional method and is able to detect lipid in the sub-picomolar range; this will facilitate analysis even from small sized samples derived from *in vivo* settings such as animal models, biopsies, micro-dissected specimens and other high value samples whose analysis might inform on PI5P metabolism or signalling. For example, over the years, it has been clear from analysis of the gene PIKFYVE in mammals, that this enzyme can synthesize PI5P [6,8,29]. However, due to its activity *in vitro* on both PI and PI3P, the route by which it synthesises PI5P *in vivo* is still debatable. The later route requires a 3-phosphatase, members of the myotubularin in mammals, to convert the PI(3,5)P_2_to PI5P. An analysis of this specific route in mammals has been limited since mammals have around 14 homologs that can potentially encode myotubularin activity [30]. *Drosophila* has only 4 predicted orthologs of myotubularin and thus offers a better metazoan model to study the existence of phosphatases that can regulate PI5P. PI5P elevations in the nucleus have been reported in the context of stress signals and DNA damaging agents such as radiation implicating this lipid in cell signalling in the setting of cancer biology [[3,31,32] and reviewed in [33]]. The availability of our highly sensitive, non-radioactive mass assay will allow the measurement of PI 5 P levels in micro-dissected biopsy samples from human tumours where the application of radioactive mass assays will be challenging. Results from such samples using our assay could help decipher signalling mechanisms in human tumours and help inform decision making in clinical oncology. Finally, PIP4K a key enzyme in the regulation of PI5P levels has been recently implicated in the control of insulin receptor signalling and Type II diabetes [23,34,35]. The mechanisms by which PIP4K regulates insulin signalling remain unclear but the use of our assay on human samples of small size and high value could help in the study of the role of this enzyme and its control of PI5P levels in human type II diabetes settings.

Although our mass assay has been developed for PI5P measurement, its use is not limited to the assay of just this lipid. In principle, it could also be used to measure the levels of other phosphoinositides that can also be assayed using an *in vitro*, enzyme based conversion mass assay [17]. Thus, the method we have described to measure PI5P is a safe and scalable technique that will be of great interest to both researchers specifically exploring PI5P metabolism and phosphoinositide signalling in general.

## Acknowledgements

We thank the NCBS Mass spectrometry facility for support.

## Funding Information

This work was funded by the National Centre for Biological Sciences-TIFR and a Wellcome DBT India Alliance Senior Fellowship to PR (IA/S/14/2/501540). VR was supported by a Research Associateship from the Department of Biotechnology, Government of India.

## Author contributions

AG, SS and DS conceptualized the project. AG, SS, VR and DS performed experiments. AG and SS analysed data. Writing: AG, SS and PR. Supervision of project: PR. Funding acquisition: PR,

## Competing interests

The authors declare that there are no competing interests associated with the manuscript.

